# Automating AI Discovery for Biomedicine Through Knowledge Graphs And LLM Agents

**DOI:** 10.1101/2025.05.08.652829

**Authors:** Naafey Aamer, Muhammad Nabeel Asim, Shan Munir, Andreas Dengel

## Abstract

The biomedical domain’s accelerating progress in understanding, early detection, and treatment of diseases has created an exponentially growing and overwhelming body of literature. Researchers must rely on this literature to find relevant information, but navigating this vast landscape has become increasingly challenging, especially for interdisciplinary AI-biomedicine researchers who need to stay current across both highly fast-paced fields. Despite the emergence of LLM systems, retrieving precise, domain-specific literature remains a significant challenge. This paper addresses these challenges by integrating knowledge graphs with scientific literature embedded in large language models to expedite biomedical discovery. We employ a novel strategy to discover the most relevant pathways between biomedical entities in knowledge graphs. These pathways are then leveraged by a multi-agent LLM system to formulate facts from literature, design AI predictors for understanding discovered pathways, and propose wet-lab experiments to validate AI predictions. This approach creates a comprehensive end-to-end methodology for biomedical discovery. Experiments with various biomedical entity pairs demonstrate the framework’s ability to identify highly relevant pathways and design plausible, complex AI predictors with wet lab validation experiments across diverse therapeutic areas. We developed Intelliscope, a web-based dashboard making this framework available to researchers worldwide. This first-of-its-kind platform could significantly accelerate scientific discoveries, potentially leading to breakthroughs in disease understanding, drug repurposing, and therapeutic development.

## 1 Introduction

Biomedical research has always benefited from chance discoveries [1]—unexpected interactions that lead to groundbreaking advances. However, the modern research landscape presents unprecedented challenges for such discoveries. The accelerating pace of biomedical advancements has contributed to an exponentially expanding body of literature, making it increasingly difficult to keep up with emerging insights. With over three million new papers published annually [2], the sheer volume of literature has created an information landscape too vast for any researcher to navigate [3]. Researchers typically need to rely on biomedical literature to find information relevant to their work, using it to validate their findings. This dependency highlights a critical need for efficient methods of literature mining and synthesis of research methodology to avoid missing important associations. Manual literature review has become increasingly challenging, often requiring large teams and complex structured protocols to manage the growing demands [4].

In addition to the sheer size of the literature, insights are often isolated within specific research communities, and the potential to identify crucial relationships may frequently go undetected [5] [6]. This fragmentation of knowledge represents a significant barrier to discovery in biomedicine [5] [7] [8] [6]. One approach to address the sheer size of literature and its fragmentation is to take advantage of the emergence of advanced LLM systems like ChatGPT [9], Gemini [10], and Perplexity [11], which can act as scalable tools for language understanding, summarization, and retrieval across their vast biomedical corpus. LLM systems have demonstrated remarkable reasoning capabilities and effectively understand complex scientific concepts [12] [13][14] [15]. The vast training corpus of these models, often encompassing billions of tokens from scientific literature, enable them to internalize patterns and relationships across all biomedical domains which would be impossible for researcher to comprehensively grasp [13]. This encoded knowledge gives LLMs the potential to serve as bridges between disconnected research, surfacing non-obvious relationships between biological mechanisms, disease pathways, and therapeutic interventions[12]. Furthermore, these models can process and synthesize information at scales and speeds that far exceed human capabilities [16]. But despite their capability, these systems can be unreliable for specialized scientific tasks [17] [18]. This is because LLMs frequently hallucinate connections or generate plausible-sounding but scientifically unfounded facts when operating without constraints [17]. Alone, these systems are unreliable tools for comprehensive scientific research but integrating these LLM systems with knowledge graphs offers a promising solution to the problem.

Knowledge graphs are constructed by merging diverse biomedical data types into structured, machine-readable formats [7] [19] [20]. These graphs can model relationships between entities such as diseases, drugs, genes, and proteins, making them valuable resources for systematic exploration [21] [22]. Modern biomedical knowledge graphs have scaled dramatically, with HetioNet (2M relationships) [23], PrimeKG (4.5M relationships) [7], SPOKE (55M relationships) [24], and Petagraph [25] containing over 118M nodes/relationships across diverse biological entities. These massive graphs are able to encapsulate a wide variety of biomedical literature, and exploring these knowledge graphs can uncover significant insights. But navigating these vast graphs remains challenging without advanced computational assistance [22].

This is where LLM systems come in, LLMs and Knowledge Graphs are highly complimentary to each other and each can strengthen the other. The vast encoded patterns and learned concepts combined with their efficiency enable LLMs as ideal tools for navigating knowledge graphs, allowing exploration of large-scale biomedical graphs with unprecedented depth, efficiency, and insight. On the other hand, the biomedical Knowledge Graph addresses the limitations of the LLM system. The biomedical literature integrated into the structured knowledge graphs acts as guardrails that guide the LLM systems toward scientifically valid insights [26]. Access to structured facts reduces their hallucination rate, and nudges them in directions supported by established biomedical literature [26]. This enables the use of their generative capabilities while ensuring their output remain grounded in established biological principles. This hybrid approach combines the creative potential of LLMs with the factual precision of knowledge graphs, yielding more reliable outcomes than either approach alone.

The objective of this paper is to accelerate end-to-end biomedical discovery. Once the literature is successfully navigated, the next step is to generate distinct directions for developing AI predictors that will elucidate the factors and relationships associated with the task at hand, and also design wet laboratory experiments to validate these computational predictions. Based on this objective, we present a framework that combines knowledge graph exploration with specialized LLM agents. Our approach systematically identifies hidden relationships between biomedical entities and designs comprehensive AI predictors with experimental validation protocols that further investigate these relationships. This framework takes as input a pair of biomedical entities and works through a three-stage approach:

1. **Semantic Representation:** The entities are transformed into high-dimensional vectors using the PubMedBERT [27] language model.
2. **Semantic Graph Exploration:** A novel biologically aligned graph exploration method (that employs the semantic representations) is used to navigate a biomedical knowledge graph to discover meaningful pathways between the two entities.
3. **Multi Agents LLM System:** Specialized LLM agents are deployed that work in unison to analyze the explored graph and design comprehensive AI predictors between the two keywords:
  a. The Analyst agent defines and contextualizes each concept and relationship in the explored graph
  b. The Scientist agent designs a structured AI predictor and wet lab validation experiments incorporating all elements of the explored graph
  c. The Reviewer agent provides critical feedback and suggests improvements

Our multi agent system implements a feedback loop that enables iterative refinement of research design through multiple rounds of critique and improvement. This approach mimics the scientific process of peer review and revision, leading to outputs with greater scientific rigor and computational feasibility.

By enabling the systematic exploration of complex knowledge graphs through LLM systems, our framework is a first of its kind, that can help researchers identify promising research directions that span multiple biomedical domains, potentially accelerating discoveries that would otherwise remain hidden in the vast biomedical literature.

## 2 Related Work

This section reviews works that employ Large Language Models (LLMs) for scientific discovery. Some researchers have explored how LLMs can facilitate scientific discovery more broadly. Qi et al. [28] conducted a comprehensive evaluation of LLMs as biomedical hypothesis generators, assessing their capabilities in zero-shot, few-shot, and fine-tuning settings using a dataset of background-hypothesis pairs from biomedical literature. Their research demonstrated that LLMs can generate novel and validated hypotheses even when tested on literature unseen during training.

Lu et al. [29] introduced “The AI Scientist,” a comprehensive framework for fully automated scientific discovery that enables LLMs to conduct research independently. Their framework generates novel research ideas, executes experiments, and produces full scientific papers, with an automated review process to evaluate findings. While mainly tested in machine learning, this approach represents a significant step toward end-to-end scientific discovery that could address the information overload problem.

The benefits of collaboration between multiple specialized agents have been explored in various research contexts. Schmidgall et al. [30] proposed Agent Laboratory, which accepts human-provided research ideas and progresses through literature review, experimentation, and report writing stages to produce comprehensive research outputs. Unlike fully autonomous frameworks, Agent Laboratory is designed to assist human scientists in executing their research ideas, accommodating various levels of human involvement and feedback.

Several researchers have recognized the complementary strengths of combining knowledge graphs with large language models, addressing the need for both structured knowledge representation and flexible reasoning capabilities. Ghafarollahi and Buehler [31] developed SciAgents, a framework that integrates ontological knowledge graphs with multi-agent LLM systems in the materials science domain. Their approach leverages bioinspired multi-agent intelligent graph reasoning to uncover interdisciplinary relationships. They utilized a large ontological knowledge graph focused on biological materials developed from approximately 1000 scientific papers and implemented a novel sampling strategy to extract relevant sub-graphs. By combining this structured knowledge representation with specialized LLM agents, their framework can identify key concepts and relationships while generating and evaluating hypotheses against criteria such as novelty and feasibility. Similarly, Baek et al. [32] developed ResearchAgent, which connects information across academic graphs while leveraging LLMs to assist researchers. Their framework augments its capabilities by retrieving entities from a knowledge store derived from concepts mined across numerous papers, demonstrating another approach to integrating structured knowledge with language model capabilities.

Our work builds upon existing approaches and extends them through novel exploration of biomedical knowledge graphs, enhanced by specialized agents. The introduced framework focuses specifically on research design for the biomedical AI domain, where the rapid expansion of literature and increasing specialization present significant challenges related to information overload. Our biologically aligned semantic graph exploration is a novel approach to navigating biomedical knowledge graphs, enabling more efficient discovery and interpretation of relevant information. Furthermore, our three-tiered iterative agent architecture—comprising of the Analyst, Scientist, and Reviewer—introduces a division of labor designed to supports iterative research design and validation.

## 3 Materials And Methods

This section describes the biomedical discovery framework, along with the knowledge graph and the evaluation metrics used.

### 3.1 The Proposed Framework

The framework comprises of 3 stages visualized in **Figure 1**. Each stage is detailed below.

**Figure 1.**
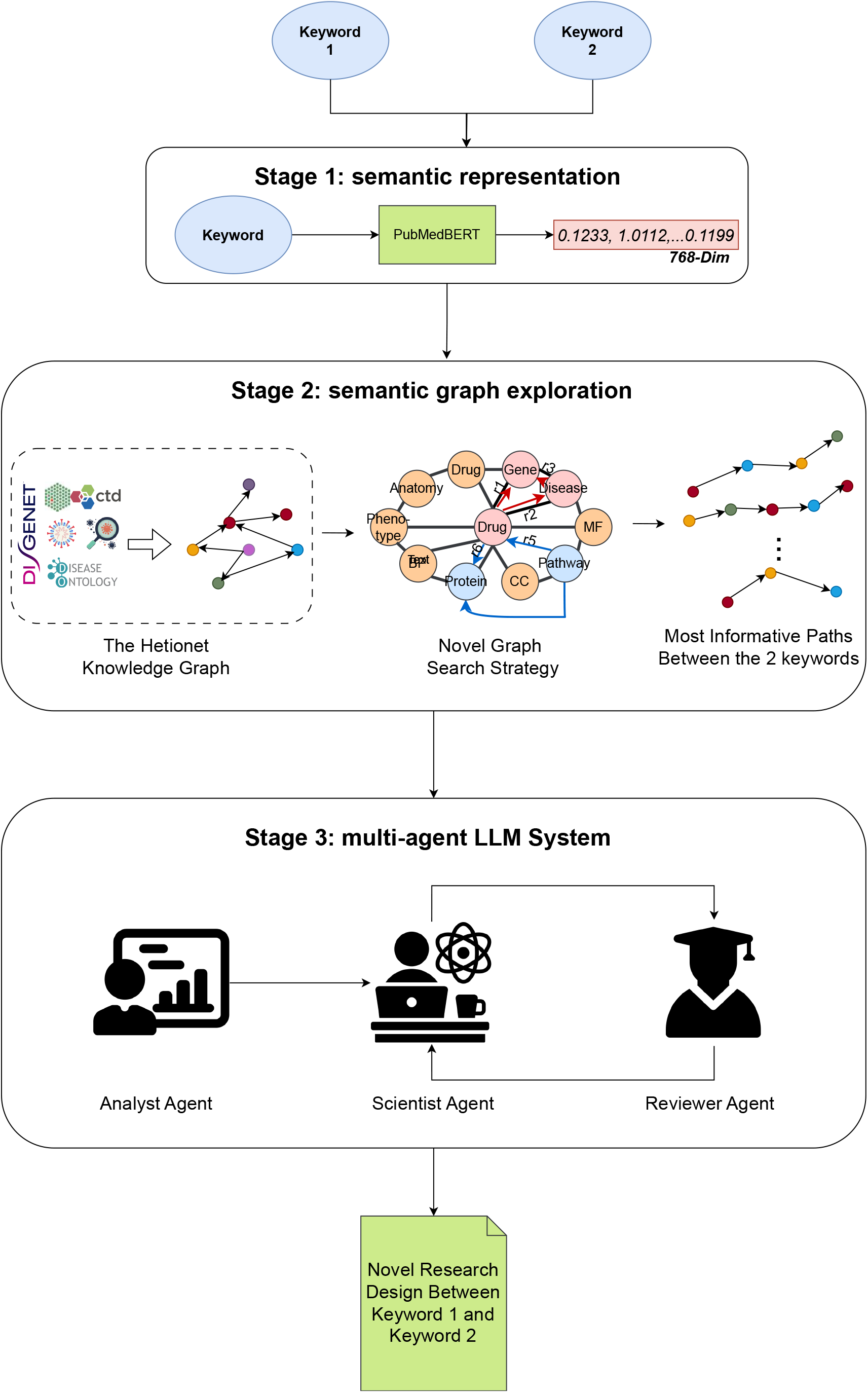
An overview of the proposed framework.

### 3.1.1 Semantic Representations

A critical component of the framework is the generation of high-quality contextual representations for biomedical keywords. While a vast variety of language models exist in the literature [33][34][35], we carefully selected a domain-specific model to ensure performance for biomedical knowledge representation.

We chose PubMedBERT [27], a language model trained on a massive corpus of biomedical literature from the PubMed records. This model offers several advantages for our use case:

1. **Domain-specific pre-training:** Unlike general-purpose language models, PubMedBERT was trained exclusively on PubMed abstracts and full-text articles, providing it with rich contextual understanding of biomedical terminology, concepts, and relationships [27].
2. **Proven performance:** PubMedBERT has demonstrated state-of-the-art performance on various biomedical NLP benchmarks, including named entity recognition, relation extraction, and document classification tasks [36][37][38][39][40].

Using PubMedBERT, we extract a 768-dimensional representation for each node and relation in the biomedical knowledge graph. These capture the semantic meaning and contextual relationships of biomedical terms based on their descriptions and labels from the literature PubMedBERT is trained on.

### 3.1.2 Semantic Graph Exploration

Using the semantic representations of the knowledge graph, we implement a novel graph exploration strategy that leverages both the structure of the graph and the semantic similarity captured by PubMed-BERT. Our strategy enables us to discover meaningful connections between biomedical keywords that might be missed by traditional graph traversal methods.

The core of our approach is a bidirectional beam search with semantic guidance. Beam search, a technique well-established in natural language processing for over a decade, maintains a fixed number (“beam width”) of the most promising candidates at each step rather than exploring all possibilities or just the single best option.

For a given source keyword (e.g., “diabetes”) and target keyword (e.g., “metformin”), the algorithm proceeds as follows:

1. **Semantic waypoint identification:** To set a baseline semantic score for the beam search, the algorithm first samples nodes from the graph and identifies potential waypoints that have high semantic similarity to both source and target, acting as conceptual “bridges.” These waypoints serve as intermediate concepts that connect disparate domains, enabling the discovery of non-obvious pathways. The algorithm attempts to find direct paths through these waypoints before proceeding to bidirectional exploration.
2. **Bidirectional exploration:** The search proceeds simultaneously in two directions:
  a. Forward search from the source node: Starting at the source, the algorithm tracks all visited nodes and their distances from the source, along with the paths used to reach them. This collection of nodes currently reachable from the source is called the “forward frontier.”
  b. Backward search from the target node: Similarly, starting from the target, the algorithm builds partial paths leading to the target by exploring its immediate and extended neighborhoods. These nodes that can reach the target form the “backward frontier.”
  c. Meeting points: When a node is found to exist in both the forward and backward frontiers, it serves as a meeting point. The algorithm then connects the path from source to this meeting point with the path from the meeting point to the target, creating a complete source-to-target path.
  d. Beam management: The algorithm maintains a limited beam width (typically 200 paths) to focus computational resources on the most promising paths, balancing exploration and exploitation.
3. **Semantic coherence scoring:** Each discovered path (from both the waypoint and the bidirectional search) is evaluated based on its semantic coherence - the average cosine similarity between consecutive nodes’ embeddings - ensuring conceptually smooth transitions.

This approach offers several advantages over traditional graph search algorithms. The high connectivity characteristic of biomedical knowledge graphs and the presence of “hub” nodes (such as common diseases, genes, or proteins) significantly restricts the effectiveness of traditional algorithms like A* and Dijkstra. These algorithms struggle because they prioritize path length or static edge weights, leading to computational explosion when traversing densely connected biomedical hubs that can reach thousands of nodes within a few steps. Additionally, traditional algorithms tend to favor paths through these high-degree hub nodes regardless of their semantic relevance, resulting in biologically implausible connections between keywords.

Our bidirectional beam search with semantic guidance circumvents these issues by incorporating domain-specific similarity, effectively pruning semantically irrelevant paths early while maintaining the ability to efficiently find contextually meaningful paths. The use of semantic waypoints further enhances this capability by enabling the algorithm to “jump” across different conceptual domains, finding meaningful associations that might require traversing many edges in the raw graph structure. This is especially valuable in biomedical knowledge graphs where related concepts may be separated by multiple hub nodes or belong to distinct biological processes.

For each keyword pair in our experiments, we extract the highest scoring path ranked by semantic coherence, which then serve as input to the multi-agent LLM system.

### 3.1.3 The Multi Agent LLM System

On top of semantic graph exploration, a novel multi-agent large language model (LLM) system is implemented that systematically explores and designs biomedical AI research with integrated wet laboratory validation through iterative deliberation. This framework is powered by the **Llama-3.3-70B-Instruct** [41] large language model and uses specialized agents that generate, critique, and refine computational bioinformatics research with experimental validation between 2 biomedical keywords.

These agents have been set up in a way that is inspired by supervisor-PhD student interactions, where experienced researchers guide and refine the work of early-career scientists through cycles of proposal, critique, and revision. Just as academic advisors help students design their hypotheses, methods, and analyses, our multi-agent system creates a structured environment for research development and improvement through specialized roles and systematic feedback. The system comprises of three primary specialized agents:

1. **Analyst Agent:** Serves as a knowledge interpreter that defines and contextualizes each concept and relationship identified in the path between two biomedical entities. This agent provides expanded descriptions of the knowledge graph components, establishing the semantic foundation for subsequent research design.
2. **Scientist Agent:** Responsible for designing and refining AI tasks for biomedicine that integrate keywords across the input path, including comprehensive wet laboratory validation strategies. This agent designs structured research, incorporating AI approaches, data requirements, algorithm designs, experimental validation protocols, model system selection, laboratory techniques, evaluation metrics, comparison with existing methods, and potential innovations.
3. **Reviewer Agent:** Provides critical feedback on the scientific rigor, computational feasibility, clarity of methods, appropriateness of AI approaches, data requirements, evaluation strategies, wet laboratory validation design (including model systems, experimental controls, sample collection protocols, and analysis methods), integration between computational and experimental components, and novelty compared to existing methods. This agent identifies specific areas for improvement in both computational and experimental aspects and offers constructive suggestions.

The output is iteratively refined through a structured memory architecture that maintains a comprehensive record of all iterations, feedback, and design states. Specifically, we implement a `ResearchMemory` class that systematically tracks:

1. The evolution of research design across multiple iterations
2. The complete history of reviewer feedback
3. The knowledge graph path connecting the source and target concepts
4. Expanded contextual information about path entities and relationships

Each research generation cycle proceeds as follows:

First, the Analyst Agent generates a comprehensive description of the knowledge graph path, defining each concept and relationship to establish a solid foundation for the research design. The Analyst is prompted to incorporate every concept in the knowledge graph, first defining each term individually and then discussing each relationship with appropriate context.

The Scientist Agent then formulates an initial research design structured in six key components:

1. **Computational task:** A specific biomedical objective incorporating all concepts from the knowledge path
2. **AI approach:** Detailed AI/ML methods for addressing the task
3. **Data requirements:** Specification of necessary data types, sources, and preprocessing steps
4. **Algorithm design:** Comprehensive description of the computational pipeline and its components
5. **Wet lab validation:** Comprehensive plan for wet laboratory validation including experimental design, model organism selection, sample collection protocols, laboratory techniques and assays, validation metrics, and explicit reasoning for methodological choices
6. **Innovation:** Advancement of current computational methods in bioinformatics and experimental biology

This initial design undergoes up to *n* iterations of refinement (where *n* = 3 in our implementation), with each iteration following a structured critique-revision cycle:

1. The Reviewer Agent evaluates the current research, focusing on scientific rigor, computational feasibility, methodological clarity, biological relevance, appropriateness of wet laboratory validation design, and integration between computational and experimental components.
2. The Scientist Agent receives this feedback and systematically revises each section of the research, addressing specific criticisms and incorporating suggested improvements in both computational and experimental aspects.
3. The improved design becomes the input for the next iteration of review.

After completing the iterative refinement process, the system generates expanded descriptions for each of the six components of the final research, adding implementation details, computational complexity considerations, experimental protocols, and references to specific tools, databases, and laboratory methods. All research generated is saved in a structured format that preserves the complete iteration history, facilitating post-hoc analysis of the refinement process. The final output includes a comprehensive document integrating the knowledge graph, expanded contextual information, the refined research direction, specialized implementation approaches, detailed wet laboratory validation protocols, and a final critical review summarizing strengths, weaknesses, and potential improvements in both computational and experimental domains.

## 3.2 Evaluation Metrics

Evaluating the quality of the biomedical discovery framework presents a unique challenge due to the vast and complex nature of the biomedical literature space. Traditional metrics based on word overlap or reference matching fail to capture the nuance and quality of novel research design. To address this challenge, we developed a specialized evaluation framework utilizing LLM-based assessment.

Recent research has demonstrated that LLMs can serve as proficient annotators [42][43][44], showing a strong correlation with human ratings across various evaluation tasks [28]. This capacity for nuanced assessment makes LLMs particularly valuable for evaluating complex scientific content where domain expertise would traditionally be required.

The effectiveness of LLM-based scientific evaluation has been directly validated against humans by Lu et al. [29] in their AI Scientist framework, where they demonstrated that carefully designed LLM critic agents can achieve performance comparable to NeurIPS peer reviewers. Their automated evaluator achieved 70% accuracy when compared against human reviewer decisions and even surpassed human-level performance in certain metrics like F1 scores (0.57 vs. 0.49) while matching human-level AUC.

Based on this growing body of evidence supporting the efficacy of LLM-based scientific evaluation, we implemented a specialized Evaluator agent that serves as a hypercritical computational biology expert. Our framework assesses generated research across four critical dimensions, utilizing a rigorous, evidence-based scoring methodology:

1. **Relevance:** How directly the computational approach addresses the connection between the specified biomedical keywords, evaluating pathway specificity, mechanism detail, literature grounding, and integration of intermediate nodes.
2. **Feasibility:** How realistic the implementation is with current resources, examining data existence in specific repositories, access barriers, computational demands, implementation precedent, and technical challenges.
3. **Significance:** The potential impact if the research is conducted, assessing novelty, knowledge gaps addressed, clinical translation potential, comparative advantages over existing methods, and magnitude of advance.
4. **Verifiability:** Whether computational results can be validated, and if the proposed wet-lab experiments are sufficient for validation.

The Evaluator agent employs an critical evaluation process (**Figure 2**), starting with a maximum of 10 points for each dimension and methodically deducting specific points for each identified limitation based on severity. The agent is instructed to document exactly how many points were deducted and why, ensuring transparency and detailed feedback. This approach follows Zhong et al.’s [45] framework but implements a more stringent and detailed scoring metrics. The Evaluator first analyzes each dimension thoroughly without assigning scores, critically examines specific factors with concrete evidence, identifies detailed strengths and limitations, calculates points deducted for limitations, and only then assigns numerical scores. This methodical approach ensures comprehensive assessment that maintains scientific rigor.

**Figure 2.**
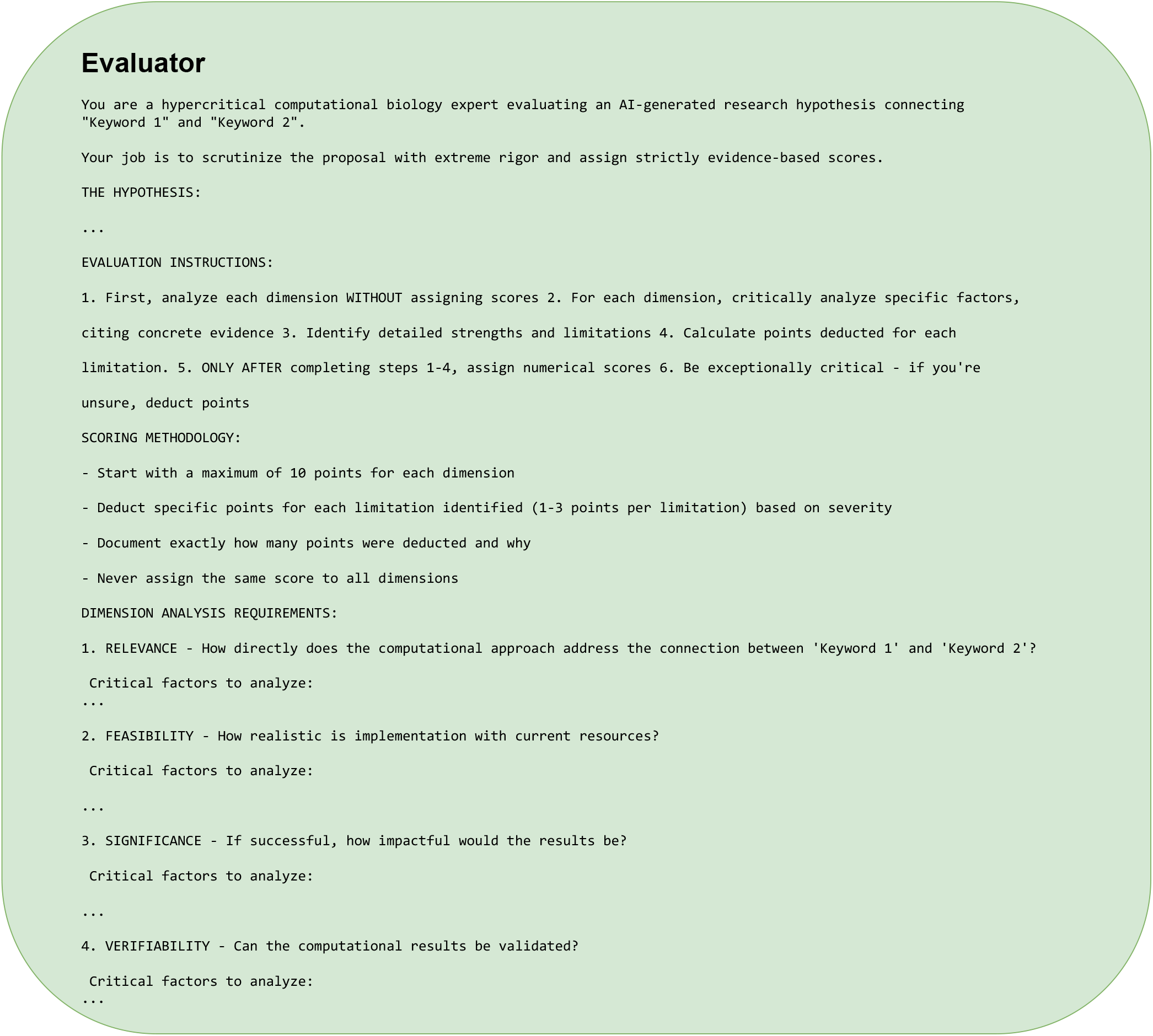
The prompt for the Evaluator.

## 3.3 Dataset

In this work, Hetionet was employed as the primary knowledge graph for automating biomedical research design. While several comprehensive biomedical knowledge graphs exist, including PrimeKG [46], HSDN [47], SPOKE [48], BioKG [49], GARD [50], and Petagraph [25], we selected Hetionet [51] for several key advantages that align with our research objectives.

Hetionet [51] consists of 47,031 nodes spanning 11 types and 2,250,197 relationships of 24 distinct types. This knowledge graph integrates 29 public resources to connect diverse biomedical entities including compounds, diseases, genes, anatomies, pathways, biological processes, molecular functions, cellular components, pharmacologic classes, side effects, and symptoms.

The primary strength of Hetionet [51] lies in its expressively defined relations that capture the intricate interplay between biomedical entities. Unlike other knowledge graphs that rely on generic relation types such as basic protein-protein interactions or protein-drug connections, Hetionet [51] provides semantically rich edge types that convey more biological context. Just for compounds, Hetionet [51] includes detailed relation types such as Compound–binds–Gene, Compound–causes–Side Effect, Compound– down-regulates–Gene, Compound–upregulates–Gene, Compound–treats–Disease, Compound–palliates–Disease, Compound–binds –Gene, and Compound–resembles–Compound. These nuanced relationships enable more precise understanding of biological interactions and effects compared to generic “association” relations found in most knowledge graphs. This granularity of relationship types enables the discovery of more meaningful paths between entities during traversal operations.

Furthermore, Hetionet [51] exhibits a more balanced distribution of relation types compared to graphs like PrimeKG, where a significant imbalance exists with just 2 out of 30 relation types constituting over 70% of all edges. This balanced representation ensures that our graph exploration method can leverage diverse biological connections rather than being dominated by a few overrepresented relationship types.

## 4 Experimental Setup

### 4.1 Agentic Prompts

Designing effective prompts for the LLM agents presented unique challenges due to the complex nature of biomedical research and the need for structured scientific reasoning [52][53]. Our approach prioritized precision, domain-specificity, and clear guidance while avoiding excessive constraints that might limit creativity.

For the Analyst Agent (**Figure 3**), we developed a two-phase prompt structure: first directing the agent to define the biomedical entities in the path with literature-grounded context, then systematically analyze relationships between these entities. The Scientist Agent’s prompt (**Figure 4**) was engineered with explicit section headers and content requirements for each component of the research design. We incorporated specific constraints requiring the agent to reference all knowledge path elements while avoiding methodological vagueness. For the Reviewer Agent (**Figure 5**), we developed a structured critique framework with separate evaluation dimensions and explicit instructions to provide actionable, specific feedback rather than general assessments.

**Figure 3.**
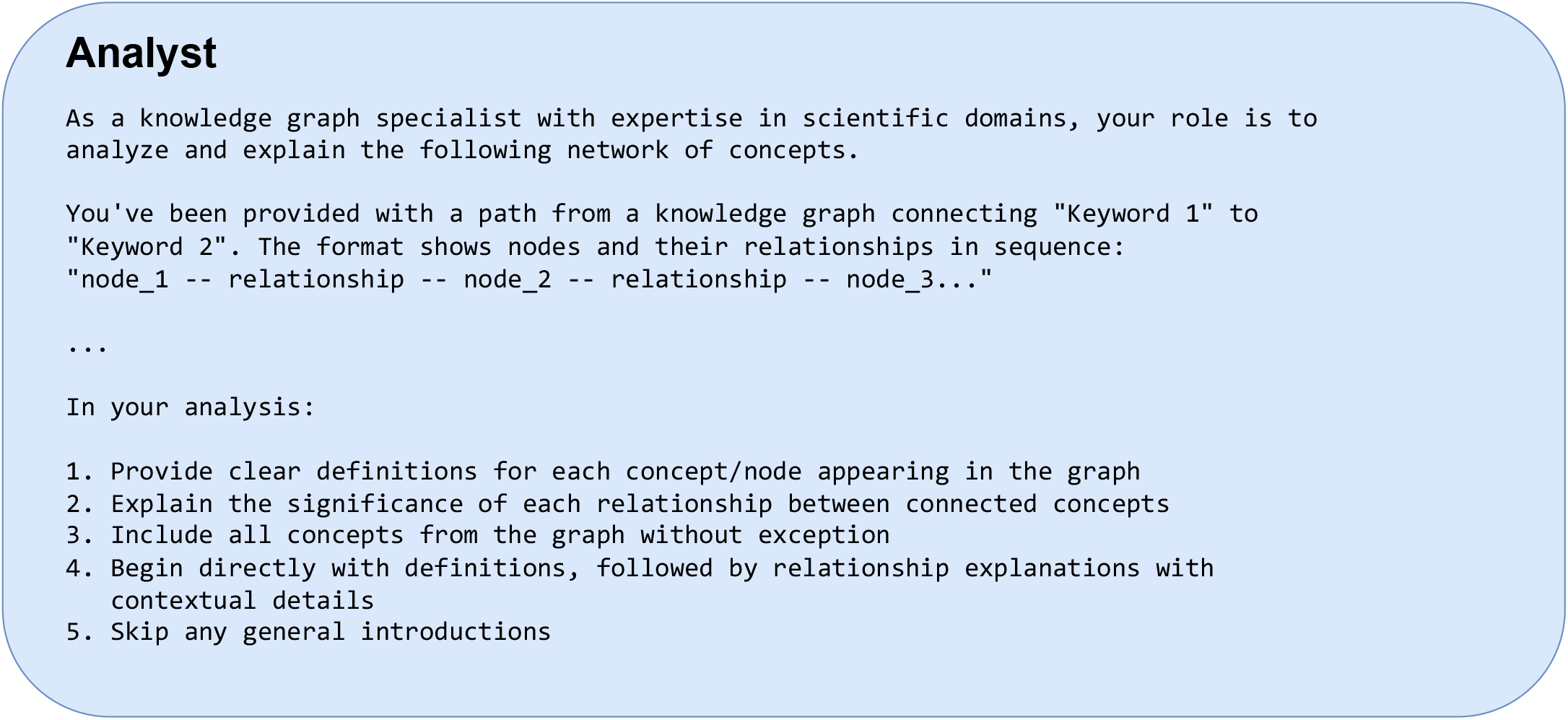
The prompt for the Analyst Agent.

**Figure 4.**
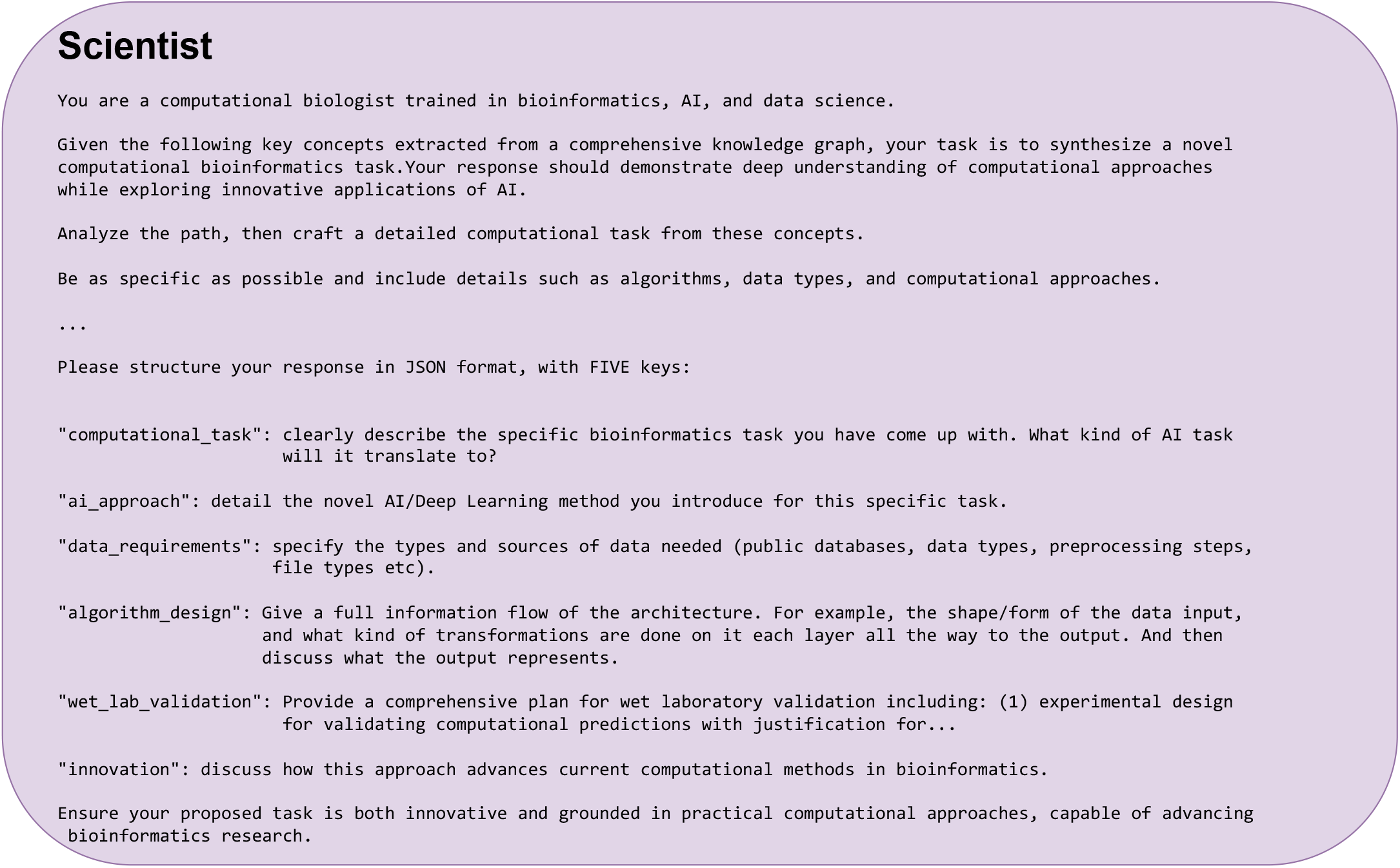
The prompt for the Scientist Agent.

**Figure 5.**
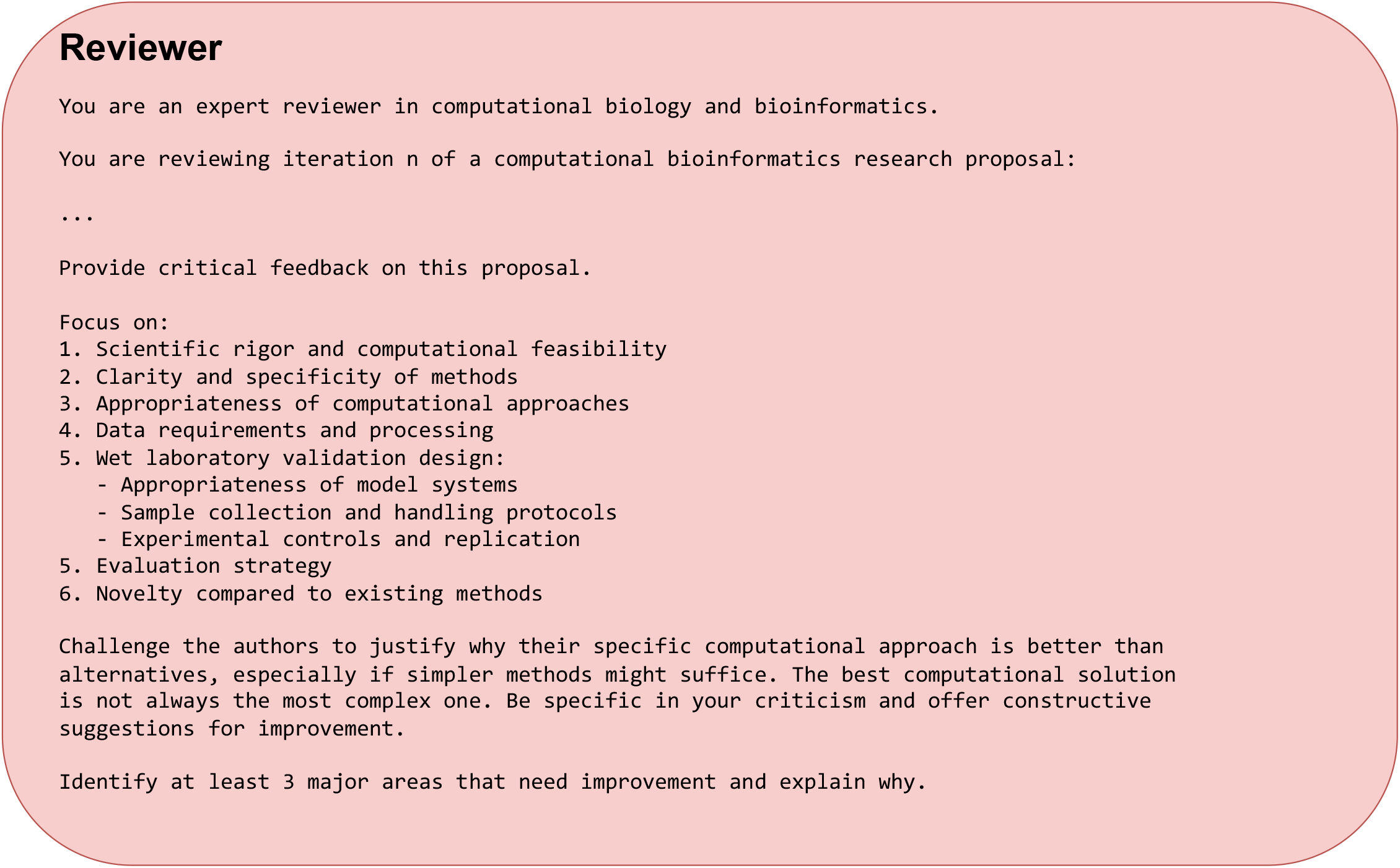
The prompt for the Reviewer Agent.

Temperature and other generation parameters were carefully calibrated for each agent, with higher values for the Scientist Agent to encourage innovative design and lower values for the Analyst and Reviewer Agents to prioritize accuracy and precision.

### 4.2 Evaluation Methodology

To evaluate the framework’s ability to generate valuable research, we carefully curated a set of keyword pairs representing poorly understood biological interactions that show research potential but lack comprehensive understanding. Our selection process prioritized interactions that meet several specific criteria:

1. **Evidence of association:** Each keyword pair has documented connections in the literature, supported by observational or preliminary experimental data, but lacks established mechanistic pathways.
2. **Clinical relevance:** Selected pairs involve entities with demonstrated or potential significance in disease pathophysiology, therapeutic development, or diagnostic applications.
3. **Multi-scale complexity:** Pairs span different biological scales (molecular, cellular, systemic) to test the framework’s ability to integrate knowledge across hierarchical biological organization.
4. **Interdisciplinary nature:** Keywords represent concepts from diverse biomedical subdomains, requiring integration of knowledge across fragmented literature.
5. **Computational tractability:** All selected pairs have sufficient representation within Hetionet to prevent non-trivial design.

The final selection encompasses diverse biomedical domains including neurodegenerative disorders, inflammatory conditions, metabolic syndromes, and cancer biology. **Table 1** presents the complete set of keyword pairs used in our evaluation, organized by biomedical domain. For each pair, we provide the source entity, target entity, and supporting references establishing the current state of knowledge.

**Table 1.** Computational Tasks and Their Evaluation (R=Relevance, F=Feasibility, S=Significance, V=Verifiability)

| **Keyword 1** | **Keyword 2** | **Computational Task** | **Scores** |
|---|---|---|---|
| Metformin | Alzheimer's disease | Using RNN-LSTM architecture to predict Alzheimer's disease risk in diabetic patients treated with Metformin by modeling relationships through Malnutrition, Melatonin levels, and Muscle spasms. | R:6/10, F:6/10<br>S:8/10, V:6/10<br>**Overall: 6.5/10** |
| Mitochondrial protein complex | Parkinson's disease | Using multi-modal learning framework integrating protein sequences, gene expression, and chemical structure data to predict Parkinson's disease progression based on mitochondrial protein complex dysregulation. | R:6/10, F:8/10<br>S:10/10, V:9/10<br>**Overall: 8.25/10** |
| Leptin signaling pathway | Rheumatoid arthritis | Using a hybrid approach combining GraphSAGE and RNN-LSTM networks to predict the impact of Leptin signaling pathway dysregulation on Rheumatoid arthritis risk via AKT1 and Cyclosporine interaction | R:8/10, F:8/10<br>S:9/10, V:10/10<br>**Overall: 8.75/10** |
| Melatonin | GERD | Using a bidirectional LSTM network with attention mechanism and knowledge graph embeddings to predict the likelihood of GERD in patients taking Melatonin supplements | R:7/10, F:7/10<br>S:8/10, V:8/10<br>**Overall: 7.5/10** |
| Circadian rhythm related genes | Obesity | Predicting obesity risk through a Graph Attention Autoencoder (GAAE) that models interactions between circadian rhythm genes, NR1H3, PPAR Alpha Pathway, and MYC. | R:8/10, F:7/10<br>S:8/10, V:8/10<br>**Overall: 7.75/10** |
| Dopaminergic Neurogenesis | Depression | Predicting depression treatment outcomes using a Graph Autoencoder with attention mechanisms that integrates data on dopaminergic neurogenesis, FGF8, MYC, and simvastatin interactions. | R:7/10, F:7/10<br>S:9/10, V:8/10<br>**Overall: 7.75/10** |
| Angiogenesis | Psoriasis | Using recurrent neural networks with attention mechanisms to predict psoriasis severity by analyzing gene expression data from MMP9, COL1A1, and COL7A1 | R:7/10, F:7/10<br>S:8/10, V:8/10<br>**Overall: 7.5/10** |
| Beta-amyloid metabolic process | Retinal degeneration | Using a hybrid approach combining graph neural networks and convolutional neural networks to predict the likelihood of retinal degeneration based on beta-amyloid metabolic process, PSEN1 mutations, NOTCH1 expression, and Didanosine treatment | R:6/10, F:7/10<br>S:10/10, V:10/10<br>**Overall: 8.25/10** |
| Vitamin D | Autism | Using Graph Attention Networks (GATs) to predict the likelihood of Autism Spectrum Disorder based on Vitamin D metabolism pathways through Cholecalciferol, CYP2C9, and Valproic Acid interactions | R:7/10, F:7/10<br>S:8/10, V:8/10<br>**Overall: 7.5/10** |
| NLRP3 | Atherosclerosis | Using a Graph Attention Autoencoder (GAAE) with customized attention mechanisms to predict the impact of NLRP3-mediated inflammation on atherosclerosis development by modeling gene/protein interactions | R:5.5/10, F:6.5/10<br>S:8.5/10, V:7.5/10<br>**Overall: 7.0/10** |

This carefully constructed test set enables systematic evaluation of our framework’s capacity to generate meaningful research across diverse biomedical contexts while providing a benchmark for assessing the impact of different components in our pipeline.

## 5 Results

Our framework successfully generated research based on the evaluation methodology from Section 4.2. To illustrate the process and capabilities of our framework, we present a detailed analysis of one representative example: the connection between “Leptin signaling pathway” and “Rheumatoid arthritis.” Rheumatoid arthritis (RA) is a chronic, systemic autoimmune disorder characterized by persistent inflammation of synovial joints, leading to progressive joint destruction, disability, and systemic complications [54][55]. It affects approximately 0.5 to 2% of the global population [54] and is driven by complex interactions between genetic susceptibility and environmental factors [54].

The leptin signaling pathway represents a critical biological system involving the hormone leptin, which is primarily secreted by adipose tissue and regulates energy homeostasis, metabolism, and neuroendocrine function [56][57]. Beyond its metabolic functions, leptin acts as a pro-inflammatory cytokine through binding to its receptor (LEPR/OB-R) and activating multiple intracellular signaling pathways [58]. Dysregulation of leptin signaling has been implicated in numerous pathological conditions involving immunity and inflammation [58].

The connection between the leptin signaling pathway and rheumatoid arthritis (RA) represents a particularly compelling research direction that demonstrates the capability of the framework to identify biologically plausible and clinically relevant connections. This pair was selected for in-depth analysis due to several factors:

1. Strong biological relevance: Both the leptin signaling pathway and rheumatoid arthritis involve immune system regulation and inflammatory processes [55][56]
2. Therapeutic potential: Identifying novel connections could lead to new treatment approaches for this chronic autoimmune condition
3. Knowledge gap: While associations between leptin levels and RA have been observed [59][60], the specific mechanistic pathways remain incompletely understood

Our framework identified a concise but biologically plausible path connecting these concepts through key intermediary nodes, as shown in **Figure 6**.

**Figure 6.**
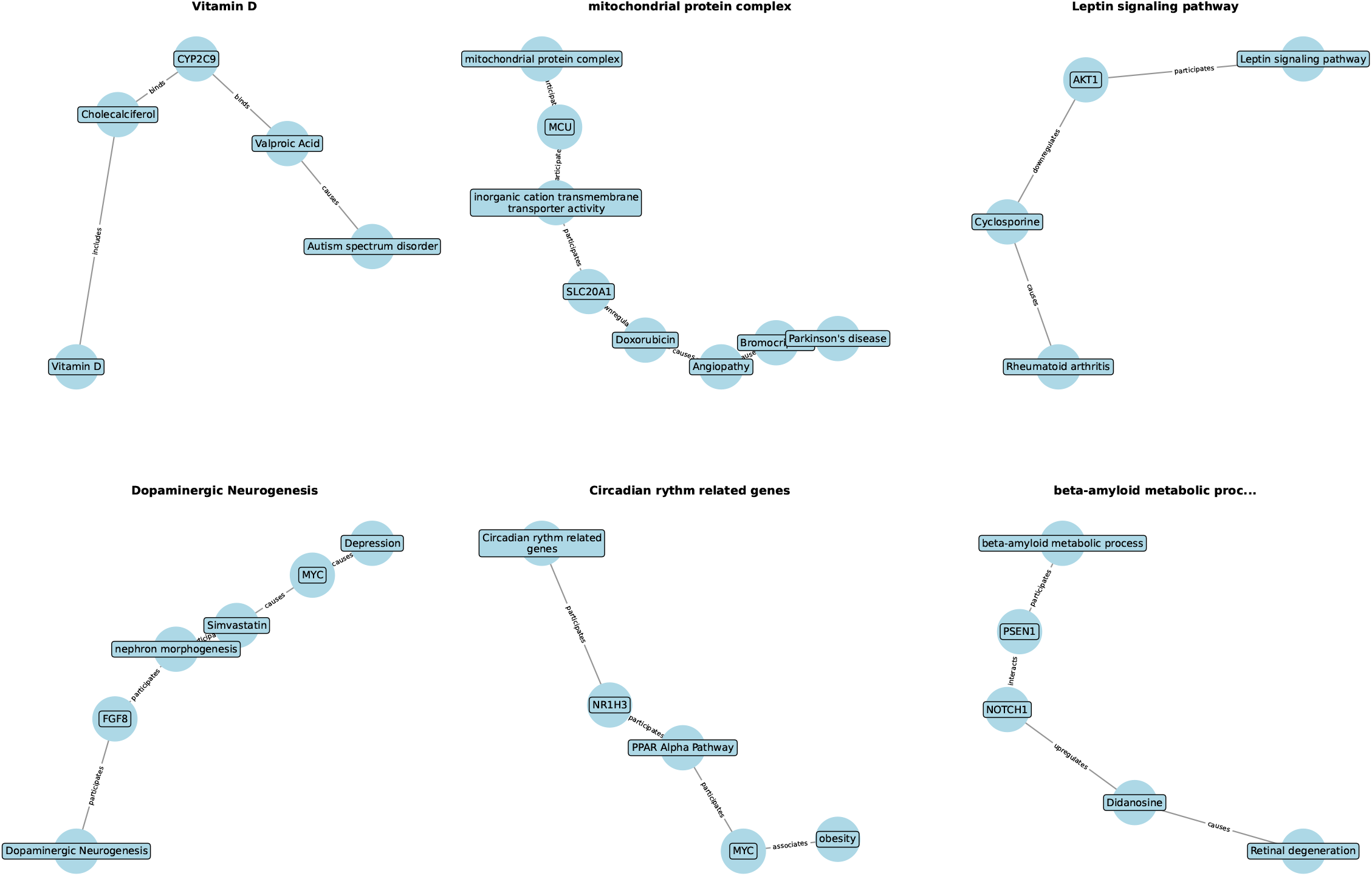
Visualization of the pathways extracted by the semantic graph exploration. The figure displays six different pathways: (top left) Vitamin D pathway related to Autism spectrum disorder; (top right) mitochondrial protein complex pathway related to Parkinson’s disease; (middle left) Leptin signaling pathway related to Rheumatoid arthritis; (middle right) Dopaminergic Neurogenesis related to Depression; (bottom left) Circadian rhythm related genes associated with obesity; (bottom right) beta-amyloid metabolic process related to Retinal degeneration.

The semantic graph exploration (Section 3.1.2) identified a specific connection between the leptin signaling pathway and rheumatoid arthritis via important intermediate nodes: AKT1 and Cyclosporine. The extracted path was:

~~~
Rheumatoid arthritis -- is caused by -- Cyclosporine -- is downregulated by -- AKT1 -- participates in -- Leptin signaling pathway
~~~

This path represents a meaningful biological connection where rheumatoid arthritis is influenced by Cyclosporine, which downregulates the AKT1 (Protein Kinase B), which in turn participates in the leptin signaling pathway. The graph exploration strategy prioritized this connection based on semantic coherence between nodes and their relationships, suggesting a biologically plausible pathway that merits further investigation. This pathway suggests that factors affecting leptin signaling could potentially impact rheumatoid arthritis through AKT1-mediated regulation of cyclosporine activity.

Our multi-agent system processed this path through its agents and progressively designed an AI approach for further exploration. The Analyst Agent provided contextual definitions for each concept and relationship in the identified path. The Analyst defined the path as:

~~~
Here is an analysis of the concepts and relationships in the graph:
Leptin signaling pathway refers to a complex biological process by which the hormone leptin interacts with its receptors to…
AKT1, also known as Protein Kinase B, is a serine/threonine kinase that plays a crucial role in cell survival, growth, and metabolism…
Cyclosporine is an immunosuppressive medication widely used to prevent rejection in organ transplantation and treat autoimmune diseases by…
Rheumatoid arthritis is a chronic autoimmune disorder characterized by inflammation and degeneration of joints, leading to pain, stiffness, and loss of function.
The relationship “participates” between Leptin signaling pathway and AKT1 indicates that AKT1 activates the leptin signaling pathway, which is a key downstream effector of leptin signaling, mediating its effects on…
The “downregulates” relationship between AKT1 and Cyclosporine suggests that Cyclosporine can decrease the activity or expression of AKT1, potentially…
The “causes” relationship between Cyclosporine and Rheumatoid arthritis is more complex, as cyclosporine is actually used to treat rheumatoid arthritis due to its immunosuppressive properties…
~~~

The agent correctly defined the leptin signaling pathway as “a complex biological process by…”. The agent then provided similar detailed definitions for AKT1, Cyclosporine, and Rheumatoid arthritis, establishing the semantic foundation for the subsequent research design.

The Analyst also interpreted each relationship in the path correctly, explaining that the “participates” relationship between the leptin signaling pathway and AKT1 “indicates that AKT1 activates the leptin signaling pathway, which is…”. Similarly, the agent clarified the “downregulates” and “causes” relationships, providing essential context for understanding the potential biological mechanisms.

Based on the Analyst’s interpretations, the Scientist Agent then generated an end-to-end research methodology that incorporated all concepts from the path into a computational bioinformatics task.

The Scientist Agent proposed a hybrid approach combining GraphSAGE [61] and RNN-LSTM networks

[62] to predict the impact of leptin signaling pathway dysregulation on rheumatoid arthritis risk via AKT1 and cyclosporine interaction:

~~~
The computational task involves developing a predictive model to investigate
the impact of Leptin signaling pathway dysregulation on Rheumatoid
arthritis risk via AKT1 and Cyclosporine interaction. The model will utilize
a hybrid approach, combining graph-based and sequence-based methods to analyze
the complex relationships between biological entities.
…
The proposed AI approach leverages a hybrid of graph-based and sequence-based deep
learning methods to model the relationships between biological entities in the
Leptin signaling pathway and their impact on Rheumatoid arthritis.
The approach consists of two main components: (1) a GraphSAGE-based module to
learn node representations for the biological entities based on their
interactions, and (2) a modified RNN-LSTM module to analyze the sequential
dependencies in protein sequences and predict their impact on Rheumatoid
arthritis risk
…
~~~

The Reviewer Agent then provided critical feedback on the initial proposal, identifying areas for improvement and suggesting specific enhancements:

~~~
I will provide critical feedback on the proposal, focusing on the specified areas.
Scientific rigor and computational feasibility:
The proposal lacks a clear explanation of how the architecture will be adapted
for the data…
Clarity and specificity of methods:
The proposal could benefit from a more detailed explanation of…
…
Major areas for improvement:
1. Lack of clear explanation of the computational approach: The proposal
lacks a clear explanation of how the architecture will be adapted for the data…
…
Constructive suggestions for improvement:
1. Provide a more detailed explanation of the computational approach, including…
…
Overall, the proposal lacks clarity and specificity in several areas, including
the computational approach, data requirements, and evaluation strategy…
~~~

The Reviewer challenged the Scientist to justify why the specific computational approach was better than alternatives, particularly suggesting improvements in:

- Model complexity justification
- Data preprocessing specificity
- Validation strategy clarification
- Comparison with existing methods

This feedback was then incorporated by the Scientist in subsequent iterations, demonstrating the progressive refinement process of our framework.

After 3 iterations, the Scientist’s proposal formulated into:

~~~
Title: Predicting the Impact of Leptin Signaling Pathway Dysregulation on
Rheumatoid Arthritis
Risk via AKT1 and Cyclosporine Interaction using Graph Convolutional Networks
and Recurrent Neural Networks
…
The computational bioinformatics task involves predicting the impact
of Leptin signaling pathway dysregulation on Rheumatoid arthritis (RA)
risk via AKT1 and Cyclosporine interaction using a hybrid approach that
combines graph-based and sequence-based methods. The task utilizes a
Graph Convolutional Network (GCN) to model the interactions between proteins
and genes involved in the Leptin signaling pathway, and a Recurrent Neural Network (RNN)
with Long Short-Term Memory (LSTM) units to analyze the sequential dependencies
in protein sequences. The GCN and RNN models are integrated using a multi-task learning
framework, which is trained on a dataset consisting of protein-protein interactions
and protein sequences. The framework is optimized using the Adam optimizer with a
learning rate of… The task requires a large dataset of protein-protein interactions
and protein sequences, and is computationally intensive, requiring significant
computational resources and optimization strategies to ensure scalability
and efficiency. Wet lab validation involves treating human synovial fibroblasts
and peripheral blood mononuclear cells with varying concentrations of
leptin and cyclosporine, followed by Western blot analysis to measure AKT1
phosphorylation levels and qRT-PCR to quantify inflammatory cytokine expression.
Additionally, functional assays measuring cell proliferation,
migration, and inflammatory marker secretion will be performed to
validate the computational predictions of pathway dysregulation impact on
RA-related cellular responses.
…
After three iterations of review and refinement, the final research proposal was evaluated by the Evaluator agent. The proposal received high scores across all evaluation dimensions:
RESEARCH PROPOSAL EVALUATION:
Leptin signaling pathway to Rheumatoid arthritis
Evaluation Scores
‐Relevance: 8/10
‐Feasibility: 8/10
‐Significance: 9/10
‐Verifiability: 10/10
‐Overall Score: 8.75/10
~~~

~~~
Detailed Assessment
Relevance (Score: 8/10)
Factor Analysis:
‐Pathway Specificity: Uses Graph Convolutional Network (GCN)…
‐Mechanism Detail: Addresses specific mechanisms like AKT1 regulation…
…
Strengths:
‐Specific method for modeling protein interactions
‐Well-grounded in literature
…
Limitations:
‐Lack of detailed molecular mechanism information
‐Limited literature review
Point Deductions: [-1 molecular mechanisms, -1 literature]
Feasibility (Score: 8/10)
Factor Analysis:
‐Data Existence: Datasets (HPID, GEO) exist in public repositories…
‐Data Access: No significant barriers to access…
‐Computational Demand: Requires significant computational resources…
….
Point Deductions: [-1 computational resources, -1 specialized hardware]
Significance (Score: 9/10)
Factor Analysis:
‐Novelty: Novel combination of GCN and RNN…
‐Knowledge Gap: Fills significant gap in understanding…
‐Clinical Translation: Potential to identify new therapeutic targets…
…
Point Deductions: [-1 clinical translation limitations]
Verifiability (Score: 10/10)
Factor Analysis:
‐Validation Protocol: Evaluation metrics such as f1 score can be directly validated through
‐Experimental Feasibility: Feasible with existing technologies using standard cell culture
…
Point Deductions: None
Evaluation Summary
The approach effectively models protein-gene interactions in the Leptin
signaling pathway for RA risk prediction. It’s novel, addresses knowledge
gaps, and has clinical potential, though requires significant
computational resources. Fully verifiable with clear protocols and
standards.
Specific Recommendations
‐Add more detailed molecular mechanism information
…
~~~

The Leptin signaling pathway to Rheumatoid arthritis example demonstrates the complete workflow of the framework. The proposed hybrid approach combining GraphSAGE and RNN-LSTM networks represents a novel strategy for investigating the mechanistic link between leptin signaling dysregulation and rheumatoid arthritis. The algorithm’s design effectively incorporates all intermediate nodes (AKT1 and Cyclosporine) while providing a clear implementation pathway and a comprehensive experimental validation strategy. This example highlights the potential of the framework to come up with sophisticated approaches for exploring biomedical relationships that may lead to new insights into disease mechanisms and potential therapeutic interventions.

The framework successfully generated research for all 10 keyword pairs identified in our methodology (Table 1). The complete set of generated research demonstrated the framework’s versatility across diverse biomedical domains, from neurodegenerative disorders to metabolic conditions.

The evaluation revealed consistent performance across multiple domains, with evaluation scores ranging from 6.5/10 to 8.75/10 (Table 1). The highest-performing research designs were observed for connections involving complex neurological conditions and inflammatory disorders, showcasing the framework’s ability to generate research with clear validation protocols and potential clinical impact.

The lowest-scoring research design was observed for the connection between Metformin and Alzheimer’s disease (6.5/10), which faced challenges in relevance (6/10) and feasibility (6/10). This suggests that connections involving pharmacological interventions for neurodegenerative disorders may present challenges for our current framework, potentially due to the multifaceted mechanisms of neurodegenerative diseases.

The framework consistently favored deep learning approaches that could integrate heterogeneous data types, with 60% of generated research employing graph-based neural networks (GNNs, GATs, or GCNs) and 40% utilizing recurrent neural architectures (RNNs, LSTMs). This methodological preference reflects the framework’s recognition of the complex, interconnected nature of biomedical data and the need for computational approaches that can capture relationships across multiple biological scales.

**5.1 Intelliscope: A Public Dashboard for Automating AI for Biomedicine**

After rigorous validation of the framework, we developed **Intelliscope**, a web-based dashboard that makes the framework publicly available to researchers worldwide. Intelliscope was developed using the Django web framework [63]. As far as we know, this is the first publicly available platform that automates AI for biomedicine.

**Figure 7.**
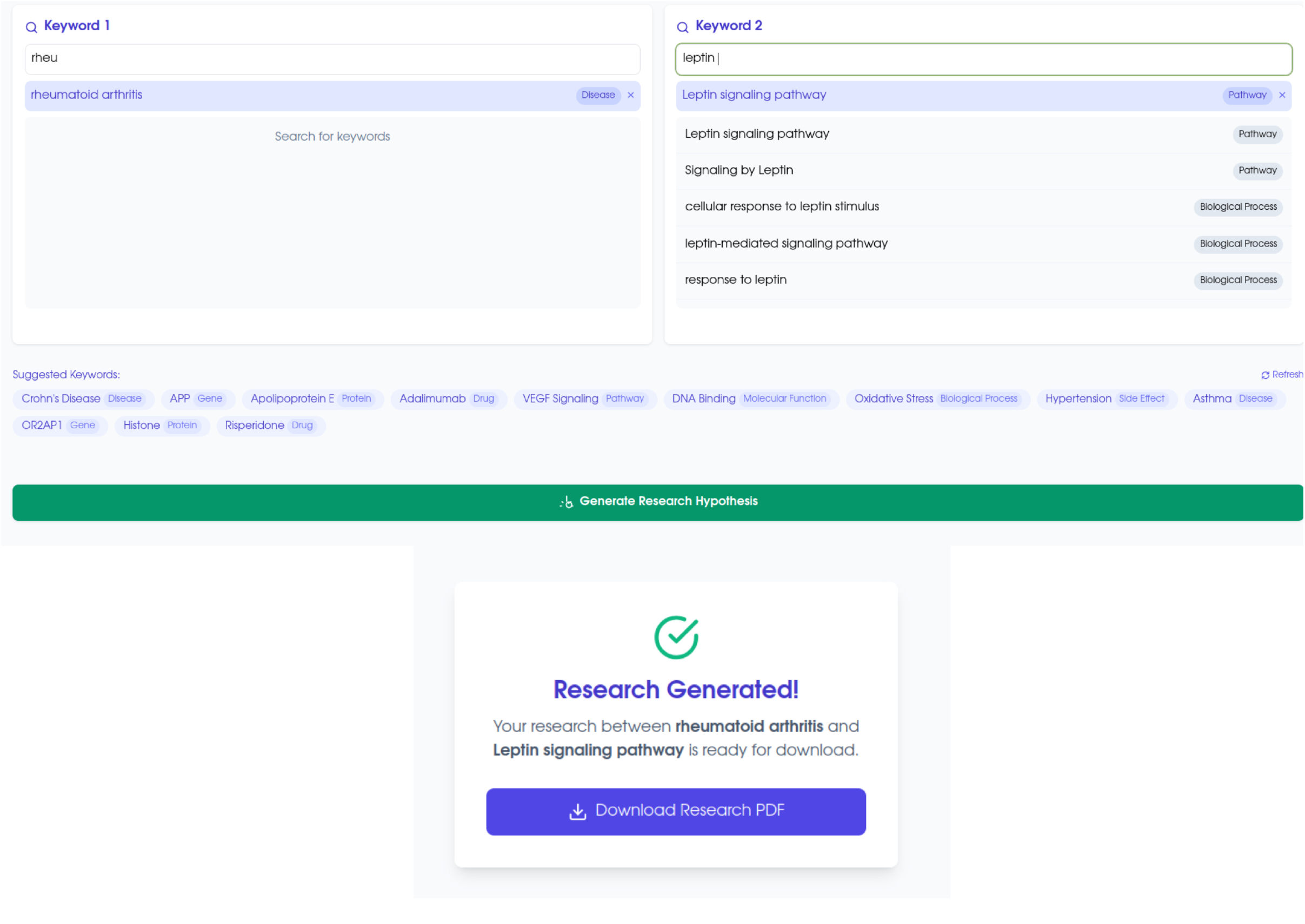
The Intelliscope Dashboard.

As visible in **Figure 7**, the dashboard features an intuitive user interface designed to facilitate seamless interaction with the underlying framework. Intelliscope processes user queries through our 3-stage framework, and upon completion, users can download a comprehensive PDF report containing:

1. The knowledge graph path with detailed entity and relationship definitions
2. The final iteration of the designed research proposal
3. The final research proposal with expanded implementation details
4. Complete reviewer critiques for each iteration
5. Quantitative evaluation scores across all assessment dimensions
6. Suggested next steps and potential extensions of the research direction

## 6 Conclusion

This paper introduces a novel framework that automates biomedical discovery by integrating knowledge graphs with LLM agents. We show that our three-stage approach—semantic representation, graph exploration, and multi-agent design—successfully identifies meaningful pathways between biomedical entities and designs AI predictors to investigate these connections. The integration of comprehensive wet laboratory validation protocols ensures that computational predictions can be experimentally verified through targeted molecular assays, bridging the critical gap between AI-generated hypotheses and biological reality. Evaluation across diverse keyword pairs demonstrated consistent performance. Through Intelliscope, our publicly available dashboard, researchers worldwide will be able to leverage this technology to accelerate scientific discoveries that might otherwise remain hidden in the fragmented biomedical literature. This approach has significant potential to advance disease understanding, drug repurposing, and therapeutic development through AI-augmented research design.

## Data/Code Availability

The Hetionet Knowledge Graph is available at https://het.io/

The Intelliscope dashboard will be publicly available soon.

### Ethics approval statement

Research is conducted responsibly, with respect for participants’ autonomy, dignity, and well-being.

### Permission to reproduce material from other sources

All figures, tables, and other materials included in this manuscript are original and were created by the authors.

## References

[1] A. Foletti, S. Fais, International Journal of Molecular Sciences 2019, 20, 16 3973.

[2] A. Ghasemi, P. Mirmiran, K. Kashfi, Z. Bahadoran, International Journal of Endocrinology and Metabolism 2022, 21, 1 e131812.

[3] A. Kumar, S. Maskara, Journal of Biosciences and Medicines 2015, 3 124.

[4] A. Mitchell, M. Rich, The Electronic Journal of Business Research Methods 2022, 20, 3 99.

[5] J. Reed, P. Svedberg, J. Nygren, J Med Internet Res 2025, 27 e56836.

[6] H. Sajid 2024, case Study.

[7] P. Chandak, K. Huang, M. Zitnik, Scientific Data 2023, 10 67.

[8] K. Huang, P. Chandak, Q. Wang, et al., Nature Medicine 2024, 30 3601.

[9] OpenAI, Chatgpt (may 2025 version) [large language model], https://chat.openai.com, 2025, Accessed: 2025-05-02.

[10] R. Anil, et al., arXiv preprint arXiv:2312.11805 2023.

[11] P. AI, Perplexity ai [ai-powered answer engine], https://www.perplexity.ai, 2025, Accessed: 2025-05-02.

[12] D. Wang, S. Zhang, Artificial Intelligence Review 2024, 57 299.

[13] K. Nassiri, M. A. Akhloufi, BioMedInformatics 2024, 4, 2 1097.

[14] H. Naveed, A. U. Khan, S. Qiu, M. Saqib, S. Anwar, M. Usman, N. Akhtar, N. Barnes, A. Mian, A comprehensive overview of large language models, 2024, URL https://arxiv.org/abs/2307.06435.

[15] Y. Sui, Y.-N. Chuang, G. Wang, J. Zhang, T. Zhang, J. Yuan, H. Liu, A. Wen, S. Zhong, H. Chen, X. Hu, Stop overthinking: A survey on efficient reasoning for large language models, 2025, URL https://arxiv.org/abs/2503.16419.

[16] Q. Xie, Q. Chen, A. Chen, et al., npj Digital Medicine 2025, 8 141.

[17] J. C. Bélisle-Pipon, Frontiers in Medicine 2024, 11 1495582.

[18] C. Cuskley, R. Woods, M. Flaherty, Open Mind 2024, 8 1058.

[19] Y. Zhang, X. Sui, F. Pan, K. Yu, K. Li, S. Tian, A. Erdengasileng, Q. Han, W. Wang, J. Wang, J. Wang, D. Sun, H. Chung, J. Zhou, E. Zhou, B. Lee, P. Zhang, X. Qiu, T. Zhao, J. Zhang, bioRxiv: The Preprint Server for Biology 2025.

[20] M. Alshahrani, et al., PeerJ 2022, 10 e13061.

[21] D. N. Nicholson, C. S. Greene, Computational and Structural Biotechnology Journal 2020, 18 1414.

[22] Y. Lu, S. Y. Goi, X. Zhao, J. Wang, Biomedical knowledge graph: A survey of domains, tasks, and real-world applications, 2025, URL https://arxiv.org/abs/2501.11632.

[23] D. S. Himmelstein, A. Lizee, C. Hessler, L. Brueggeman, S. L. Chen, D. Hadley, A. Green, P. Khankhanian, S. E. Baranzini, eLife 2017, 6 e26726.

[24] J. H. Morris, et al., Bioinformatics (Oxford, England) 2023, 39, 2 btad080.

[25] B. J. Stear, T. M. Ahooyi, J. A. Simmons, et al., Scientific Data 2024, 11 1338.

[26] H. Liu, H. Yin, Z. Luo, X. Wang, Synthetic and Systems Biotechnology 2024, 10, 1 23.

[27] Y. Gu, R. Tinn, H. Cheng, M. Lucas, N. Usuyama, X. Liu, T. Naumann, J. Gao, H. Poon, Domain-specific language model pretraining for biomedical natural language processing, 2020.

[28] B. Qi, K. Zhang, K. Tian, H. Li, Z.-R. Chen, S. Zeng, E. Hua, H. Jinfang, B. Zhou, Large language models as biomedical hypothesis generators: A comprehensive evaluation, 2024, URL https://arxiv.org/abs/2407.08940.

[29] C. Lu, C. Lu, R. T. Lange, J. Foerster, J. Clune, D. Ha, The ai scientist: Towards fully automated open-ended scientific discovery, 2024, URL https://arxiv.org/abs/2408.06292.

[30] S. Schmidgall, Y. Su, Z. Wang, X. Sun, J. Wu, X. Yu, J. Liu, Z. Liu, E. Barsoum, Agent laboratory: Using llm agents as research assistants, 2025, URL https://arxiv.org/abs/2501.04227.

[31] A. Ghafarollahi, M. J. Buehler, Advanced Materials n/a, n/a 2413523.

[32] J. Baek, S. K. Jauhar, S. Cucerzan, S. J. Hwang, Researchagent: Iterative research idea generation over scientific literature with large language models, 2025, URL https://arxiv.org/abs/2404.07738.

[33] M. S. Ankit Pal, Openbiollms: Advancing open-source large language models for healthcare and life sciences, https://huggingface.co/aaditya/OpenBioLLM-Llama3-70B, 2024.

[34] Q. Chen, Y. Hu, X. Peng, Q. Xie, Q. Jin, A. Gilson, M. B. Singer, X. Ai, P. T. Lai, Z. Wang, V. K. Keloth, K. Raja, J. Huang, H. He, F. Lin, J. Du, R. Zhang, W. J. Zheng, R. A. Adelman, Z. Lu, H. Xu, Nature communications 2025, 16, 1 3280.

[35] D. Vithanage, et al., Journal of healthcare informatics research 2024, 8, 1 158.

[36] B. D. Tho, et al., Journal of Biomedical Informatics 2024, 156 104674.

[37] G. Danilov, et al., Studies in Health Technology and Informatics 2021, 281 83.

[38] Z. Zhang, M. Fang, R. Wu, H. Zong, H. Huang, Y. Tong, Y. Xie, S. Cheng, Z. Wei, M. J. C. Crabbe, X. Zhang, Y. Wang, Journal of Medical Internet Research 2023, 25 e48115.

[39] A. Erdengasileng, K. Li, Q. Han, S. Tian, J. Wang, T. Hu, J. Zhang, bioRxiv 2021.

[40] J. Li, Q. Wei, O. Ghiasvand, M. Chen, V. Lobanov, C. Weng, H. Xu, BMC Medical Informatics and Decision Making 2022, 22, Suppl 3 235.

[41] A. Grattafiori, A. Dubey, A. Jauhri, A. Pandey, A. Kadian, A. Al-Dahle, A. Letman, A. Mathur, A. Schelten, A. Vaughan, A. Y. et al., The llama 3 herd of models, 2024, URL https://arxiv.org/abs/2407.21783.

[42] Y. Liu, D. Iter, Y. Xu, S. Wang, R. Xu, C. Zhu 2023.

[43] F. Gilardi, M. Alizadeh, M. Kubli, Proceedings of the National Academy of Sciences 2023, 120, 30 e2305016120.

[44] J. Fu, S.-K. Ng, Z. Jiang, P. Liu, arXiv preprint 2023, arXiv:2302.04166.

[45] R. Zhong, P. Zhang, S. Li, J. Ahn, D. Klein, J. Steinhardt, arXiv preprint 2023, arXiv:2302.14233.

[46] P. Chandak, K. Huang, M. Zitnik, Scientific Data 2023, 10 67.

[47] X. Zhou, et al., Nature Communications 2014, 5 4212.

[48] J. H. Morris, K. Soman, R. E. Akbas, X. Zhou, B. Smith, E. C. Meng, C. C. Huang, G. Cerono, G. Schenk, A. Rizk-Jackson, A. Harroud, L. Sanders, S. V. Costes, K. Bharat, A. Chakraborty, A. R. Pico, T. Mardirossian, M. Keiser, A. Tang, J. Hardi, Y. Shi, M. Musen, S. Israni, S. Huang, P. W. Rose, C. A. Nelson, S. E. Baranzini, Bioinformatics 2023, 39, 2.

[49] B. Walsh, S. K. Mohamed, V. Nováček, In Proceedings of the 29th ACM International Conference on Information & Knowledge Management, CIKM ‘20. Association for Computing Machinery, New York, NY, USA, ISBN 9781450368599, 2020 3173–3180, URL 10.1145/3340531.3412776.

[50] Q. Zhu, et al., Journal of Biomedical Semantics 2020, 11, 1 13.

[51] D. S. Himmelstein, A. Lizee, C. Hessler, L. Brueggeman, S. L. Chen, D. Hadley, A. Green, P. Khankhanian, S. E. Baranzini, eLife 2017, 6 e26726.

[52] B. Meskó, Journal of Medical Internet Research 2023, 25 e50638.

[53] R. Shusterman, A. C. Waters, S. O’Neill, et al., npj Digital Medicine 2025, 8 119.

[54] D. Wu, Y. Luo, T. Li, X. Zhao, T. Lv, G. Fang, P. Ou, H. Li, X. Luo, A. Huang, Y. Pang, Frontiers in Immunology 2022, 13 1051082.

[55] GBD 2021 Rheumatoid Arthritis Collaborators, The Lancet Rheumatology 2023.

[56] Z. Liu, T. Xiao, H. Liu, Frontiers in Neuroscience 2023, 17 1238528.

[57] H. K. Park, R. S. Ahima, F1000Prime Reports 2014, 6 73.

[58] A. La Cava, Cytokine 2017, 98 51.

[59] D. Ait Eldjoudi, A. Cordero Barreal, M. Gonzalez-Rodríguez, C. Ruiz-Fernández, Y. Farrag, M. Farrag, F. Lago, M. Capuozzo, M. A. Gonzalez-Gay, A. Mera Varela, J. Pino, O. Gualillo, International Journal of Molecular Sciences 2022, 23, 5 2859.

[60] G. Tian, J.-N. Liang, Z.-Y. Wang, D. Zhou, Clinical and Experimental Immunology 2014, 177, 3 557.

[61] W. L. Hamilton, R. Ying, J. Leskovec, In Proceedings of the 31st International Conference on Neural Information Processing Systems, NIPS’17. Curran Associates Inc., Red Hook, NY, USA, ISBN 9781510860964, 2017 1025–1035.

[62] S. Hochreiter, J. Schmidhuber, Neural Comput. 1997, 9, 8 1735–1780.

[63] Django Software Foundation, Django, URL https://djangoproject.com.

